# Differential SP1 Interactions in SV40 Chromatin from Virions and Minichromosomes

**DOI:** 10.1101/2020.05.17.100925

**Authors:** Kincaid Rowbotham, Jacob Haugen, Barry Milavetz

## Abstract

SP1 binding in SV40 chromatin in vitro and in vivo was characterized in order to better understand its role during the initiation of early transcription. We observed that chromatin from disrupted virions, but not minichromosomes, was efficiently bound by HIS-tagged SP1 *in vitro*, while the opposite was true for the presence of endogenous SP1 introduced in vivo. Using ChIP-Seq to compare the location of SP1 to nucleosomes carrying modified histones, we found that SP1 could occupy its whole binding site in virion chromatin but only the early side of its binding site in most of the minichromosomes carrying modified histones due to the presence of overlapping nucleosomes. The results suggest that during the initiation of an SV40 infection, SP1 binds to an open region in SV40 virion chromatin but quickly triggers chromatin reorganization and its own removal by a hit and run mechanism.

## INTRODUCTION

The regulation of eukaryotic gene expression is thought to occur primarily through the binding of transcription factors to the cognate DNA sequences present in the chromatin of the gene to be regulated. The binding of the transcription factors then serves as a catalyst for the addition of other factors and subsequent reorganization of the pre-existing chromatin structure. While this general mechanism is particularly relevant to cellular regulation, it is also thought to represent the events occurring in viral DNA that is organized into chromatin during the course of an infection.

The polyomaviruses, of which Simian Virus 40 (SV40) is a member, are a family of small circular DNA viruses that exist as chromatin throughout their life cycle and as consequence should be subjected to regulation in the same way as cellular chromatin. For SV40 there is evidence that this is true. SV40 is regulated by many of the same transcription factors binding to their cognate DNA that function in eukaryotic cells (Tooze, 1981; Wasylyk et al., 1983; Wildeman, 1988) and SV40 chromatin contains the same chromatin structural elements found in cellular chromatin including a nucleosome-free regulatory region, specifically positioned nucleosomes, and histone modifications (Balakrishnan and Milavetz, 2017a, b; Kube and Milavetz, 1989, 1996; Kumar et al., 2017; Kumar et al., 2018; Scott and Wigmore, 1978; Varshavsky et al., 1979).

During the course of an SV40 infection, there is a time dependent shift from early transcription to late transcription and this shift is associated with changes in the organization of nucleosomes and histone modifications in the viral chromatin (Balakrishnan et al., 2010; Balakrishnan and Milavetz, 2005; Kallestad et al., 2014; Kallestad et al., 2013; Kumar et al., 2017). Because the SV40 genome is organized as chromatin in virions, the chromatin present in the virions serves as a potential link between the chromatin structure present late in infection that serves as the substrate for encapsidation and the chromatin structure that is required to initiate a subsequent infection when the virion infects a cell. In order to test whether the formation of virions was associated with changes in chromatin structure, we compared the chromatin structure of SV40 minichromosomes late in infection (the putative encapsidation substrates) to the structure of the chromatin found in infectious virions (Kumar et al., 2018).

We observed that in both form of SV40 chromatin a nucleosome containing various histone modifications was present in the SV40 enhancer region in a large proportion of the chromatin with one important difference. In minichromosomes isolated late in infection the nucleosome was located over the early side of the enhancer and extended into the SP1 binding sites, while in the chromatin from virions, the nucleosome appeared to have shifted to the right into the late region and exposed the early side of the enhancer and all of the SP1 binding sites (Kumar et al., 2018). Because the location of this nucleosome would be expected to affect the binding of transcription factors that regulate early and late transcription (Tooze, 1981; Wasylyk et al., 1983; Wildeman, 1988), we hypothesized that the sliding nucleosome would result in a shift from a chromatin structure that favored late transcription in the substrates for encapsidation to a structure that favored early transcription in the chromatin from virions (Kumar et al., 2018).

If this hypothesis is correct, we would expect to see differences in the binding of transcription factors to the SV40 regulatory region in the different forms of SV40 chromatin. In order to test this, we have investigated the binding of the transcription factor SP1 to the SV40 regulatory region in chromatin. We chose to use SP1 for this analysis for a number of reasons. First, SP1 is a critical transcription factor for SV40 (Dynan and Tjian, 1983; Wildeman, 1988). Second, the binding of SP1 is likely to differ between early and late transcription since early transcription requires all of the SP1 binding sites to be available (Wildeman, 1988). Third, SP1 binds with relatively high affinity (Letovsky and Dynan, 1989) which we expect would keep the SP1, if present, bound to at least some of the chromatin during our various analyses. SP1 binding was analyzed in vitro by measuring the binding of HIS-tagged SP1 to the different forms of SV40 chromatin. SP1 binding was also analyzed in vivo using ChIP and ChIP-Seq procedures to determine the amount and position of the SP1 in SV40 chromatin isolated from infected cells for comparison to the location of nucleosomes and histone modifications. These studies indicate that HIS-tagged SP1 can bind to chromatin from SV40 virions but not intracellular minichromosomes and that the failure to bind to minichromosomes is likely a result of both nucleosome positioning and the presence of SP1 bound *in vivo*.

## RESULTS

### HIS-tagged SP1 binds efficiently to chromatin from disrupted virions but not intracellular minichromosomes isolated 30 minutes or 48 hours post-infection

If the location of a nucleosome within the SV40 enhancer affected the accessibility of the SP1 binding sites located in the regulatory region as proposed (Kumar et al., 2018), we would expect that the binding of SP1 to SV40 chromatin from disrupted virions would be greater than from other forms of SV40 chromatin based upon our ChIP-Seq results. In order to determine whether the SP1 binding sites were affected by the location of the enhancer nucleosome, we measured the ability of HIS-tagged SP1 to bind to different forms of chromatin using a HIS-tagged protein pulldown assay. The results of this analysis for chromatin from disrupted SV40 virions, minichromosomes isolated 30 minutes post-infection, and minichromosomes isolated 48 hours post-infection are shown in Figure 1. As shown in the figure we observed an approximately 250-fold increase in the binding of HIS-tagged SP1 to chromatin from disrupted virions (approximately 3.5% bound) compared to the chromatin isolated at 30 minutes and 48 hours post-infection (essentially no binding). The fold difference was determined by comparing the amount bound by HIS-tagged SP1 to a similar aliquot of chromatin without the SP1.

**Figure 1.**
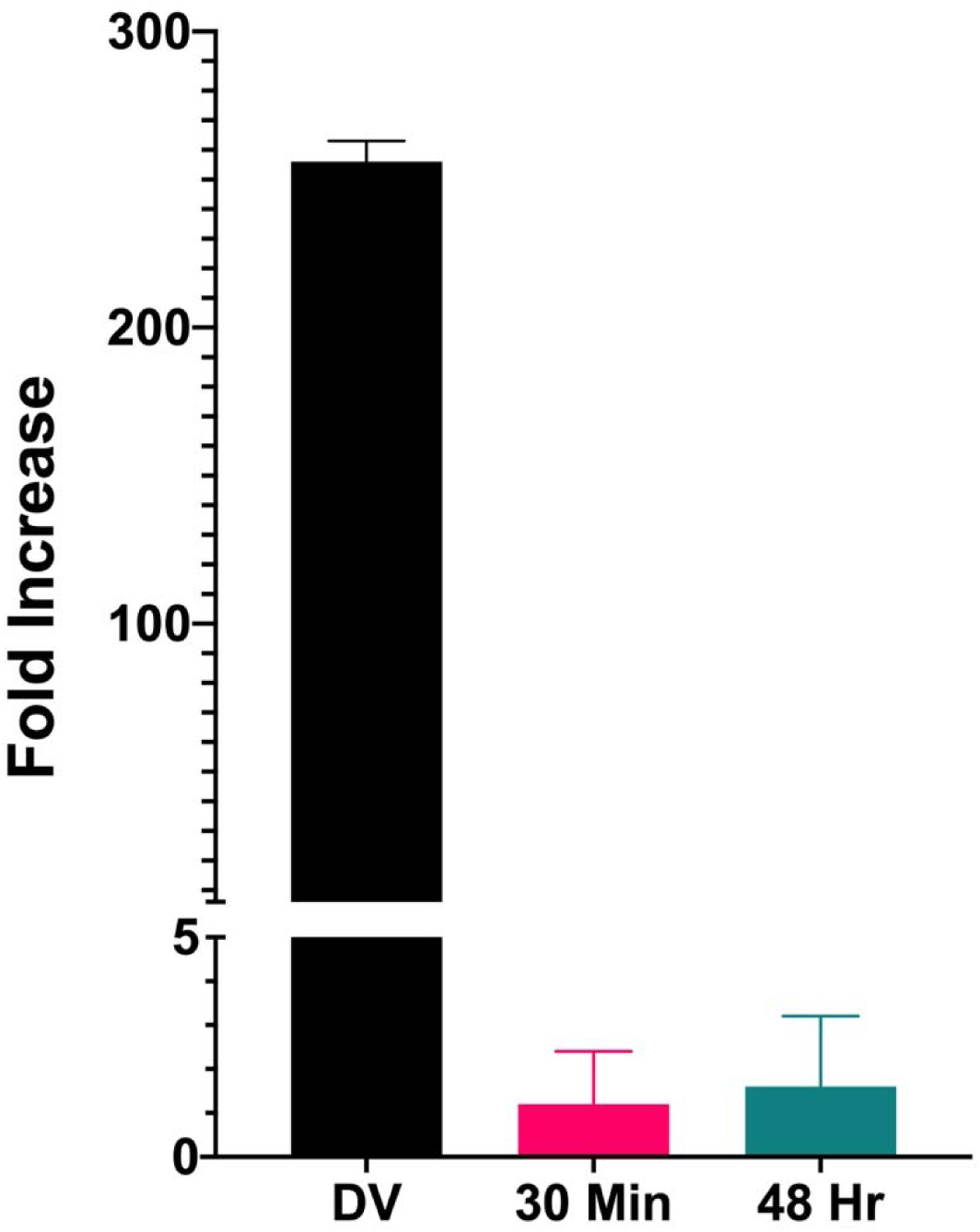
Binding of HIS-tagged SP1 to SV40 chromatin from disrupted virions and minichromosomes isolated 30 minutes and 48 hours post infection. Similar amounts of chromatin from disrupted virions, minichromosomes isolated 30 minutes post infection, and minichromosomes isolated 48 hours post infection were incubated in parallel with HIS-tagged SP1. Bound chromatin was separated from unbound chromatin and purified on Ni magnetic beads. The DNA present in the bound chromatin was purified and amplified by real-time qPCR using primers recognizing the enhancer region of the SV40 DNA. The results are displayed as the fold increase in binding in the presence of HIS-tagged SPI compared to a parallel sample treated the same without HIS-tagged SP1 being present. For each type of SV40 chromatin, at least four biological replicates were averaged to generate the graphs in the figure.

If the binding of HIS-tagged SP1 was related to the relative openness of the regulatory region in chromatin from disrupted virions, we would expect that in chromatin from the SV40 mutant cs1085 whose regulatory region has been shown to be much more open by a number of criteria (Kube and Milavetz, 1989, 1996; Kumar et al., 2017) there should be greater binding by SP1. As shown in Figure 2, there is approximately a seven-fold increase in binding to chromatin from disrupted cs1085 virions compared to disrupted wild-type virions. Taken together these results indicate that the SP1 binding sites are more accessible in chromatin from disrupted virions than other forms of SV40 chromatin and that the binding is related to the “openness” of the chromatin within the regulatory region.

**Figure 2.**
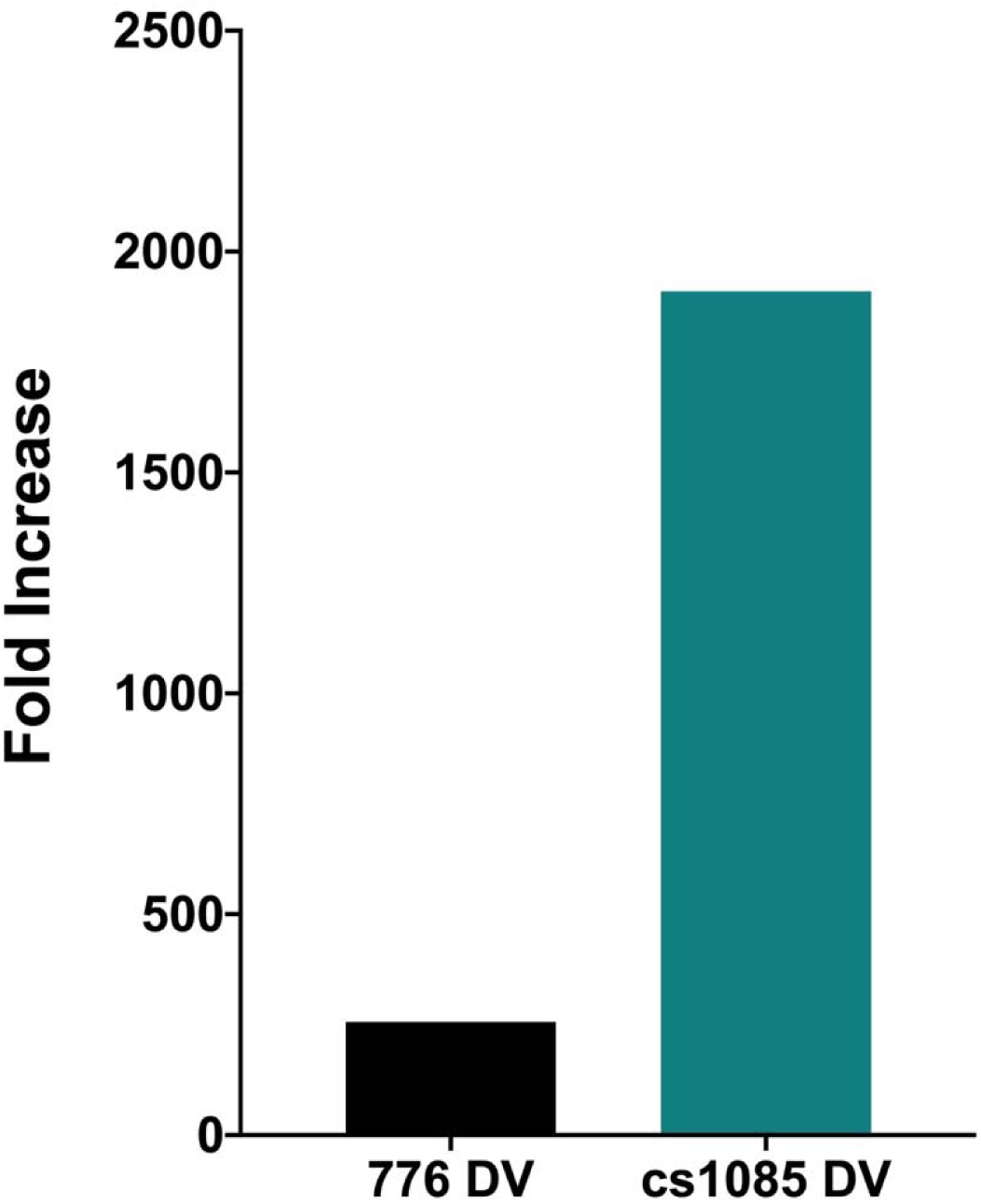
Binding of HIS-tagged SP1 to wild type and cs1085 chromatin from disrupted virions. Similar amounts of chromatin from disrupted virions prepared from SV40 wild type and the mutant cs1085 virus were incubated in parallel with HIS-tagged SP1. Bound chromatin was separated from unbound chromatin and purified on Ni magnetic beads. The DNA present in the bound chromatin was purified and amplified by real-time qPCR using primers recognizing the enhancer region of the SV40 DNA. The results are displayed as the fold increase in binding in the presence of HIS-tagged SPI compared to a parallel sample treated the same without HIS-tagged SP1 being present. For each type of SV40 chromatin, at least four biological replicates were averaged to generate the graphs in the figure.

### ChIP analysis of SP1 present in 776 DV, 48 hours and 30 minutes

The 50-fold increase in binding by HIS-tagged SP1 of chromatin from disrupted wild-type virions compared to other chromatin suggested that the binding site for SP1 was open in the virion chromatin. In order to test whether this was the case we analyzed the SV40 chromatin for the presence of SP1 loaded in vivo using standard ChIPs. Chromatin was incubated with antibody to SP1 and the percentage of the input chromatin that was bound was determined by real-time PCR. Results of this analysis are shown in Figure 3. We observed approximately a 45-fold reduction (compared to the 48 hour minichromosomes) in the percentage of chromatin from disrupted SV40 virions which contained SP1 (0.1±0.1% of input) compared to the 30 minute (1.0±0.2%) and 48 hour minichromosomes (4.5±3.5% of input). The results from this analysis are complementary to those with the HIS-tagged SP1 and indicate that the chromatin from virions contains very little if any SP1 bound and consequently the SP1 binding sites are available for binding by the transcription factor. In contrast, the two forms of minichromosomes appear to contain similar relative amounts of SP1 within the regulatory region of the genome when isolated from infected cells.

**Figure 3.**
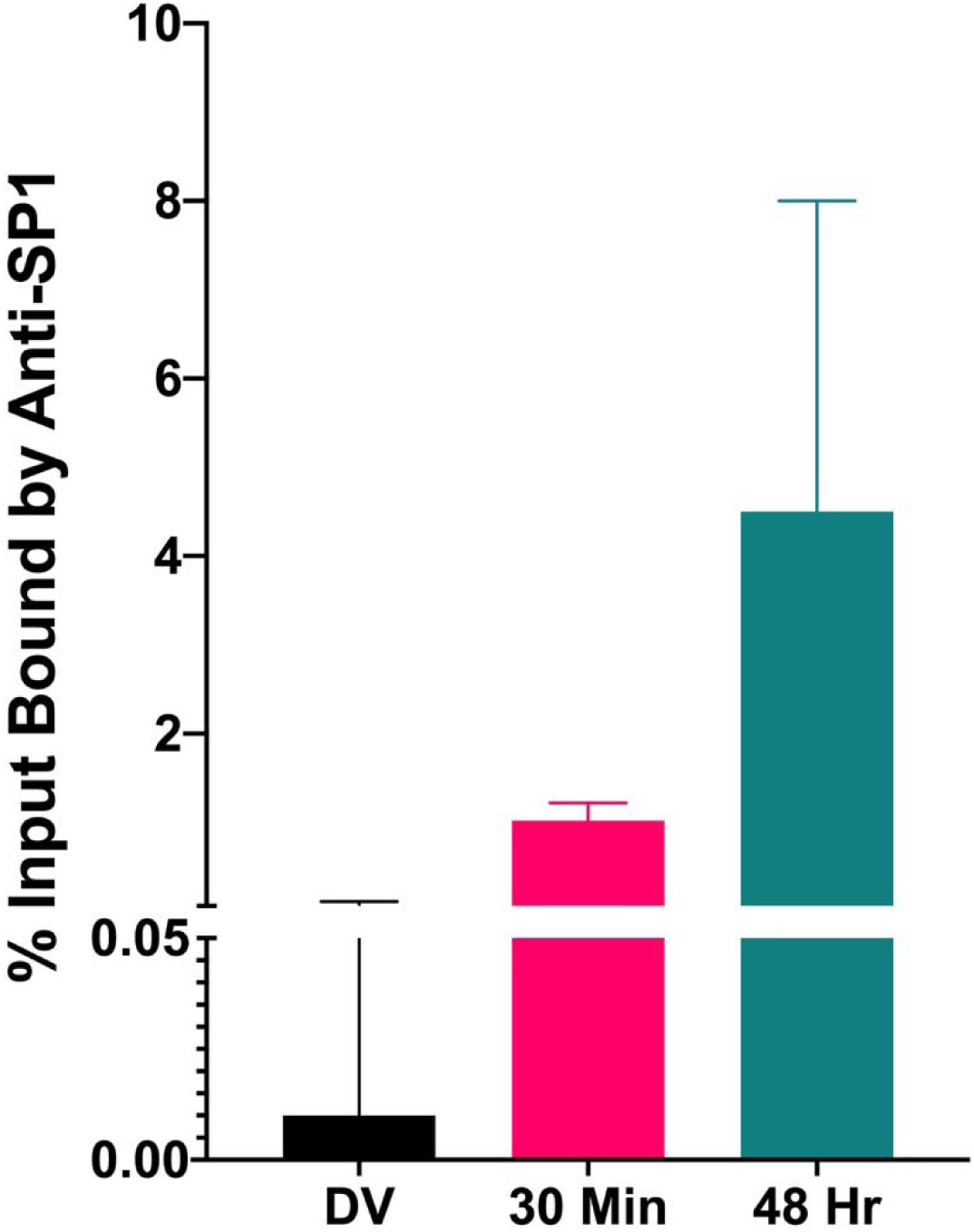
Presence of endogenous SP1 in SV40 chromatin from disrupted virions and minichromosomes isolated 30 minutes and 48 hours post infection. Similar amounts of chromatin from disrupted virions and minichromosomes isolated 30 minutes and 48 hours post infection were subjected to a ChIP assay with antibody to SP1. The DNA present in the bound chromatin was purified and analyzed along with an aliquot of the corresponding input chromatin by real-time qPCR using primers that recognize the enhancer region. The percentage of input chromatin that was bound by SP1 was calculated for each sample. For each type of SV40 chromatin, at least four biological replicates were averaged to generate the graphs in the figure.

### Organization of SP1 in SV40 chromatin

Since SP1 appeared to be present in a significant fraction of the SV40 minichromosomes isolated at 30 minutes and 48 hours post infection, we determined the location of the SP1 in the regulatory region of the chromatin from these time points. For a number of reasons, it seemed likely that the binding of SP1 in the various forms of SV40 chromatin would differ with respect to the occupancy of the GC binding sites. First, it has been shown that binding of SP1 to activate early transcription requires the whole GC binding region that consists of six tandem copies of the sequence GGGCGG separated by two to four bases consisting of either A, T, or C (Wildeman, 1988), suggesting that at 30 minutes post-infection all six GC sites will be occupied by SP1. In contrast, late transcription has not been shown to have the same requirement so that it is possible that SP1 would be bound differently in the 48 hour minichromosomes. Second, the location of nucleosomes within the regulatory region in chromatin from virions, 30 minute minichromosomes, and 48 hour minichromosomes also vary significantly with the virions and 30 minute minichromosomes more (open) than the 48 hour minichromosomes (Kumar et al., 2017; Kumar et al., 2018).

In order to test whether the binding of SP1 differed in the various forms of SV40 chromatin, we prepared sequencing libraries from SP1 ChIPs of SV40 minichromosomes isolated at the two times of interest. Following sequencing and bioinformatics analysis of the data, heatmaps were prepared by merging at least 4 biologic replicates for each sample and the location of SP1 in the two forms of chromatin compared to the location of nucleosomes determined by using the FS kit from New England Biolabs (*manuscript submitted*) and the location of nucleosomes containing HH3, HH4, H3K9me1, and H3K9me3. The latter four histone modifications were chosen because they consist of modifications that are thought to be associated with activated chromatin (HH3 and HH4) and repressed chromatin (H3K9me1 and H3K9me3) and are present in relatively high percentages of minichromosomes. In addition, the locations of these histone modifications in the regulatory region are characteristic of all the histone modifications tested to date. Because SP1 would be expected to protect approximately 60 base pairs of DNA when it bound all the GC sites in its cognate sequence, the location of SP1 was determined using sequencing reads from 75 to 99 base pairs in length. The results of this comparison are shown in Figure 4.

**Figure 4.**
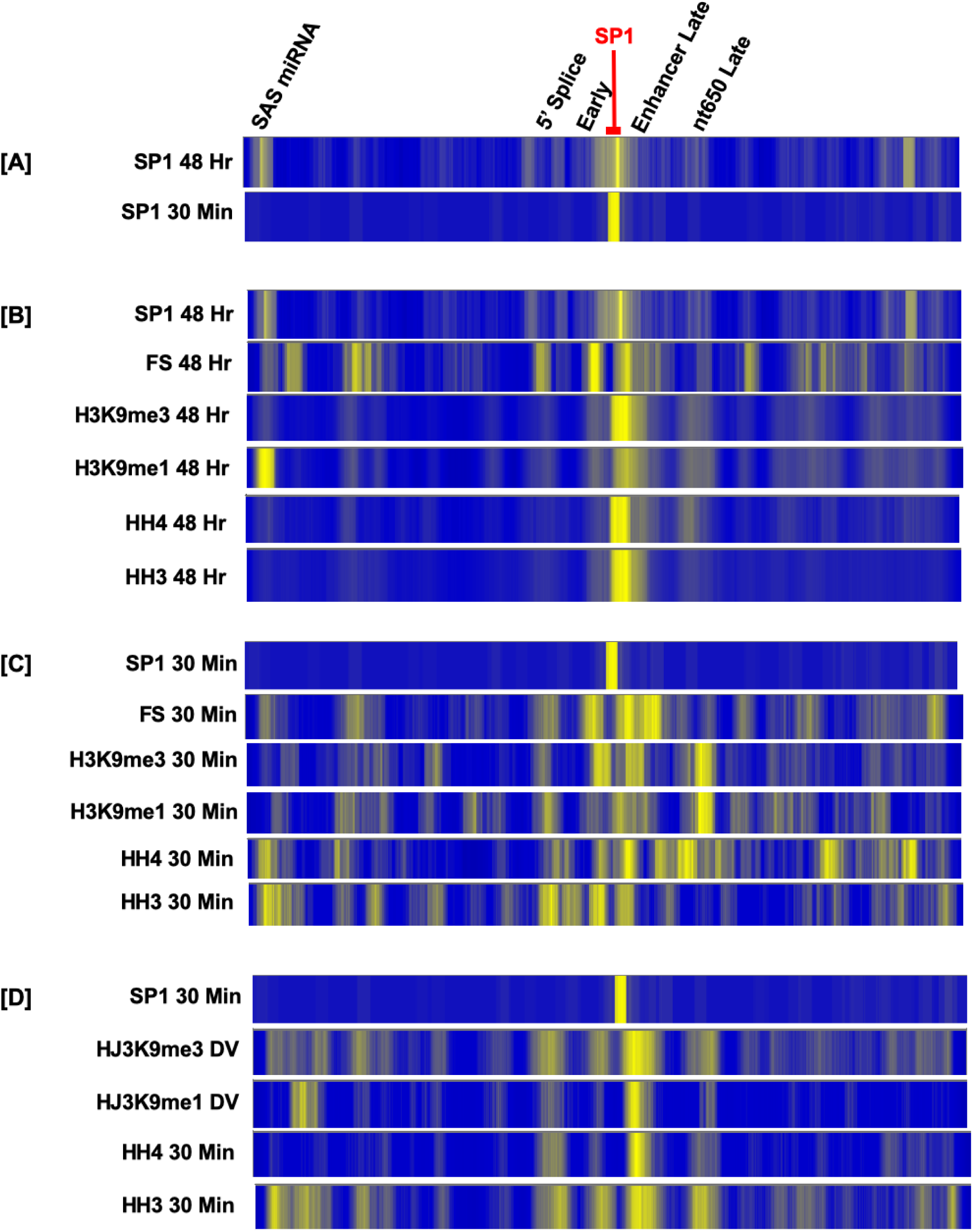
Location of SP1 in minichromosomes isolated 30 minutes and 48 hours post infection in comparison to nucleosomes carrying HH3, HH4, H3K9me1, and H3K9me3 in SV40 chromatin from minichromosomes isolated 30 minutes post infection, 48 hours post infection and disrupted virions. Similar amounts of chromatin from disrupted virions and minichromosomes isolated 30 minutes and 48 hours post infection were subjected to ChIP assays with antibodies to SP1, HH3, HH4, H3K9me1, or H3K9me3. Following washing according to the protocol found in the Millipore-Sigma kit, the chromatin bound by each antibody was fragmented by sonication and the bound chromatin separated from the other chromatin fragments. The DNA present in each bound fraction was purified and used for library preparation using the NEB Next UltraII kit. Libraries prepared with each antibody were sequenced on a MiSeq and analyzed using bioinformatics tools to determine the location of the DNA present in each library on the SV40 genome. For the SP! DNA libraries, reads from 75 to 99 base pairs in size were analyzed to correspond to the approximate size of the SV40 DNA protected by SP1. For the antibodies recognizing histone modifications, reads from 100 to 150 base pairs in size were analyzed since they correspond to the approximate size of the DNA present in a nucleosome. For each combination of SV40 chromatin and antibody, at least four biological replicates were merged to generate the heatmaps sh the appropriately sized reads are located at a site in the SV40 genome. The location of major landmarks on the genome including the enhancer, SP1 binding sites, early start, late start, early 5’ splice, nt650, and the SAS/miRNA are all indicated.

Comparing the location of the brightest band in the regulatory region, we found that SP1 appeared to be located primarily on the late side of the binding site at 48 hours and more or less over the complete GC binding site at 30 minutes, Figure 4A. In the regulatory region of SV40 minichromosomes isolated at 48 hours post infection the location of nucleosomes using FS mapping and the location of nucleosomes containing HH3, HH4, and H3K9me1 were relatively similar with major bands over the enhancer, the early RNAPII binding site, and the late side of the enhancer. Comparing the location of nucleosomes mapped with the FS kit to the location of SP1, it appeared that the first three or four GC sites from the early side in the SP1 binding region appeared to be open while the last two or three appeared to be covered by a nucleosome in a fraction of the minichromosomes. This organization was also seen in nucleosomes carrying HH3, HH4, H3K9me1, and H3K9me3 where the bound SP1 from the ChIP analysis appeared to be located primarily over the last two or three copies of the GC sites. However, in a fraction of the minichromosomes at this time the whole SP1 binding site appeared to be occupied by nucleosomes carrying these histone modifications. The results in minichromosomes isolated 48 hours post infection suggest that SP1 is present in a subset of minichromosomes primarily over the late side of the SP1 binding region.

The relationship between SP1 and nucleosome organization in SV40 minichromosomes isolated 30 minutes post infection shares some similarities and some differences with the results obtained with minichromosomes isolated 48 hours post infection. The organization of nucleosomes by FS mapping was very similar to the results obtained from minichromosomes isolated 48 hours post infection, although there were differences in the intensity of the bands. The organization of nucleosomes containing the studied modified histones showed much more variability over the region bound by the SP1. HH3, HH4, and H3K9me3 appeared to have an open region of varying size over GC sites I to III but not IV to VI in all of the minichromosomes. H3K9me1 appeared to substantially cover all six GC sites.

Since the organization of nucleosomes carrying HH3, HH4, H3K9me1, and H3K9me3 indicated that SP1 was not able to bind all six GC sites in a large fraction of minichromosomes isolated 30 minutes post infection, we compared the binding of SP1 at this time to the organization of nucleosomes carrying HH3, HH4, or H3K9me1 in chromatin from disrupted virions which would be expected to serve as the initial substrates for SP1 binding (Figure 4D). As shown in the figure the region bound by SP1 at 30 minutes would be completely accessible in the virion chromatin containing HH4 and H3K9me1 and partially accessible in the chromatin containing HH3 and H3K9me3. The change in nucleosome organization carrying these histone modifications between the chromatin in virions and the 30 minute minichromosomes indicated that there has been extensive chromatin reorganization by 30 minutes post infection with nucleosomes containing all four histone modifications moving into the GC sites from both sides of the region.

### Identification of histone modifications present in chromatin from virions preferentially bound by HIS-tagged SP1

The chromatin found in disrupted SV40 virions has been shown to be relatively heterogeneous with respect to histone modifications (Milavetz et al., 2012) and nucleosome organization (Kumar et al., 2017; Kumar et al., 2018). Since certain chromatin structural features like acetylation and methylation on H3K9 are mutually exclusive, we have proposed that there are multiple forms of SV40 chromatin present in virions and in minichromosomes (Milavetz et al., 2012). Moreover, an analysis of nucleosome positioning in chromatin from virions containing different forms of histone modifications suggested that the size of the region available for binding to SP1 differed depending upon the form of histone modification present in the chromatin (Figure 4D).

The presence of multiple forms of chromatin in virions that contain subtle differences in the size of the regulatory region potentially able to bind SP1 raised the interesting question whether SP1 binds the different forms of chromatin similarly. In order to determine whether HIS-tagged SP1 can bind to chromatin containing different histone modifications with the same affinity, we have used a two-step strategy. In the first step we used HIS-tagged SP1 to pulldown chromatin from SV40 virions. Following the pulldown, the bound SV40 chromatin was sonicated and the bound fraction containing the HIS-tagged SP1 was separated from the unbound chromatin fragments that would be expected to consist of chromatin fragments originating from the rest of the genome. In the second step of the strategy, the unbound chromatin fragments were then subjected to a ChIP analysis using antibodies to either HH3, HH4, H3K9me1 or H3K9me3. Typically, in the ChIP portion of these analyses, equal aliquots of the released fragmented chromatin from the HIS-tagged SP1 pulldown were analyzed in parallel ChIPs. The DNA present in the bound chromatin from each ChIP analysis was purified and subsequently analyzed by real-time PCR.

Since we always observed the greatest amount of DNA from the ChIP sample prepared with antibody to H3K9me1 we set the amount in each set of parallel experiments to 100% and show the relative amount of each of the other ChIPs samples as a percentage of the H3K9me1 amount in Figure 5. As shown in this figure the results indicated that HIS-tagged SP1 bound chromatin containing either H3K9me1, H3K9me3, HH3, or HH4 with relatively similar affinity although there was a clear preference for the chromatin containing H3K9me1. The results indicated that neither the form of histone modification in the chromatin or the relative size of the SP1 binding region had a major effect on the binding of SP1.

**Figure 5.**
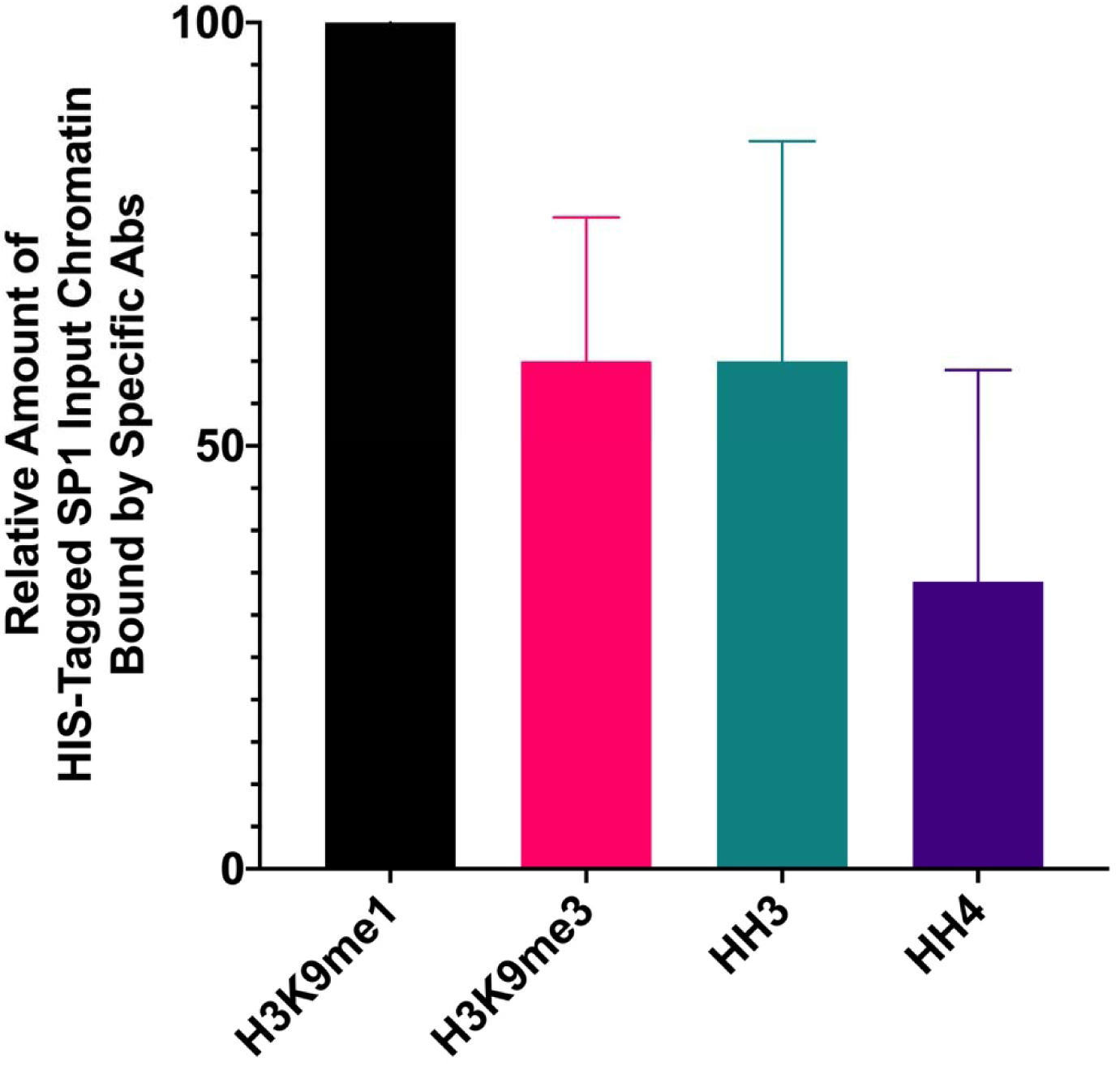
Presence of HH3, HH4, H3K9me1, and H3K9me3 in chromatin from disrupted virions bound by HIS-tagged SP1. Chromatin from SV40 disrupted virions was incubated with HIS-tagged SP1 and bound to Ni magnetic beads. Following washing to remove unbound chromatin, the Ni bound chromatin was fragmented by sonication and the bound fraction separated from the unbound chromatin fragments. The unbound fragments of SV40 chromatin were subjected to ChIP assays in parallel with antibodies to HH3, HH4, H3K9me1, and H3K9me3. The DNA present in the fractions bound by each of the antibodies were quantitated by real-time qPCR along with an aliquot of the input unbound fragments of chromatin using primers that recognize the enhancer region of the SV40 genome. For each antibody the percentage of the input fragments was calculated from the input and bound fractions. Since the amount of fragmented chromatin bound by H3K9me1 was always greatest, its amount was set as 100 for the graph. The relative amount of binding for each of the other antibodies is displayed as a percentage of the amount of product generated by the H3K9me1. For each antibody used, at least four biological replicates were averaged to generate the graphs in the figure.

## DISCUSSION

The regulation of SV40 transcription by SP1 binding to the six GC sites in the control region has been extensively studied in vivo and in vitro with similar results (Wildeman, 1988). For maximal early transcription the complete region from site I to site VI is required, but interestingly GC site IV which binds SP1 weakly is not needed (Wildeman, 1988). When the GC sites were individually mutated to reduce SP1 binding, the amount of early transcription was reduced but not equivalently for each GC site. Mutation of sites I to III affect what is referred to as the early-early RNAPII binding site while mutation of sites IV to VI affect the late-early RNAPII binding site (Wildeman, 1988). The early-early and late-early refer to the start sites for early transcription that are present either early in an infection or at times when late transcription is also occurring, respectively. The results from our ChIP-Seq analyses with antibody to SP1 were consistent with this early-early and early-late shift. We observed SP1 over the entire six GC sites at 30 minutes post infection when early-early transcription is expected, and primarily over sites V and VI at 48 hours post infection when late-early transcription is expected.

The organization of the SV40 GC sites into chromatin adds another level of regulatory complexity, since a GC site associated with a nucleosome is not bound well by SP1 (Li et al., 1994). The results that we obtained support this observation. The chromatin from disrupted virions that we previously showed to be nucleosome-free over the six GC sites (Kumar et al., 2018) was efficiently bound by HIS-tagged SP1, while the chromatin in minichromosomes isolated 30 minutes and 48 hours post infection was not. Since the HIS-tagged SP1 appeared to bind chromatin from disrupted virions similarly regardless of the histone modifications present in the chromatin, it would seem that the binding is primarily related to the fact that the GC sites are available.

The inability of HIS-tagged SP1 to bind to the chromatin from minichromosomes was not simply due to the location of nucleosomes, since we observed that both of these forms of SV40 chromatin contained SP1 using ChIP assays. As expected SP1 from a previous infection did not appear to be present in the chromatin from virions. The presence of SP1 in minichromosomes isolated 30 minutes and 48 hours post infection was not unexpected, since it has been shown to be involved in the regulation of early and late transcription. However, what was surprising was that the location of nucleosomes in minichromosomes isolated 30 minutes post infection did not appear to favor SP1 binding over all six of the GC sites. Based upon the location of nucleosomes with or without histone modifications, it appeared that by 30 minutes post infection nucleosome reorganization in the infecting viral chromatin had occurred in a substantial fraction of minichromosomes and the resulting sliding of nucleosomes was responsible for limiting the binding of SP1. By 30 minutes post infection only GC sites 1 to IV appeared to remain accessible in most of the minichromosomes, while sites V and VI appeared to contain a nucleosome in many of the minichromosomes. In minichromosomes containing H3K9me1, it appeared that reorganization resulted in nucleosomes completely blocking the GC sites. The rapid reorganization of nucleosomes containing various histone modifications could be a direct result of hit and run binding by SP1, since SP1 is known to interact with a number of chromatin modifiers including p300 and SWI/SNF (Beishline and Azizkhan-Clifford, 2015). Of course, other regulatory factors may also be involved in the reorganization process, since the enhancer is located adjacent to the SP1 region. For example, an AP1 site is located at the left end of the enhancer adjacent to the SP1 region (Wildeman, 1988) which is also likely to be exposed in the chromatin from virions.

Assuming that SP1 is at least in part responsible for driving the reorganization of nucleosomes, it is apparently acting on incoming viral chromatin regardless of the histone modifications present in the region surrounding its binding site. Since the reorganization of nucleosomes containing these different histone modifications also differ with respect to their positioning in the regulatory region after reorganization, it would appear that the nature of the histone modifications in a regulatory region of target chromatin plays an active role in determining the chromatin-related outcome upon binding by a transcription factor.

The presence of HH3 and HH4 in chromatin is generally associated with active biological processes like transcription and both histone modifications have been found to be associated with SV40 transcription (Balakrishnan and Milavetz, 2005, 2006, 2007a, b, 2008). Since nucleosomes containing HH3 and HH4 are rapidly reorganized and slide into the region occupied by SP1, it seems unlikely that SP1 remains bound to the minichromosomes containing HH3 and HH4 that have been extensively reorganized. This result suggests that SP1 is acting by a “hit and run” regulatory mechanism. Regulation of transcription by a hit and run mechanism was proposed a number of years ago (Horikoshi et al., 1988; Schaffner, 1988) and evidence supporting this idea has been obtained more recently (McNally et al., 2000) (Para et al., 2014; Varala et al., 2015). Whether SP1 functions by a hit and run mechanism during the initiation of early transcription will require further investigation.

## MATERIALS AND METHODS

### Cells and Viruses

BSC-1 cells (ATCC, CCL-26) were used in all SV40 studies including the preparation of stocks for infections, the preparation of minichromosomes, and the preparation of virion chromatin. The wild-type 776 and mutant cs1085 SV40 viruses used were gifts from Dr. Daniel Nathans. We have previously described in detail the conditions utilized for cell culture (Balakrishnan and Milavetz, 2017b).

### Infections and Purification of Minichromosomes

The procedures that we use for the preparation of minichromosomes have been described in detail (Balakrishnan and Milavetz, 2017b). In brief, SV40 minichromosomes, were prepared from sub-confluent monolayers of BSC-1 cells infected with 50 PFU per cell of a working stock of SV40 virus originally prepared at low multiplicity of infection. After 48 hours of incubation nuclei were prepared from the infected cells by washing with non-ionic detergent in a low-ionic strength buffer. The nuclei were extracted, and then purified by sedimentation in 1.4 ml Eppendorf tubes containing 10% glycerol in low-ionic strength buffer. Aliquots were collected from the top of the tube and fractions 3-5 which contain SV40 minichromosomes were pooled for subsequent analyses.

### Purification of virions and chromatin from virions

A detailed description of the procedures used for the preparation of chromatin from SV40 virions has been described in detail (Balakrishnan and Milavetz, 2017b; Kumar et al., 2017). Briefly, chromatin was prepared from low-multiplicity stock virus in order to limit the potential presence of defective virus. The stock virus was frozen and thawed at least twice to break up cells and disaggregate the virus. The SV40 virus was first concentrated by centrifugation at 50,000 x g for 35 minutes which pellets the virus and large cellular debris. The pelleted virus was resuspended in a low-ionic strength Tris-EDTA buffer and digested at least three times with DNase I at 37° C to remove any cellular or viral DNA which might be associated with the outside of the virus. Following the third digestion step, an aliquot was removed and analyzed by submerged agarose gel electrophoresis to determine whether there was any non-SV40 DNA present. Typically, three treatments with DNase I was sufficient to remove any external DNA. The nuclease-treated virus was then pelleted through 10% glycerol in low-ionic strength buffer at 50,000 x g for 35 minutes to remove any contaminants freed by the nuclease treatment and to again concentrate the virus. The nuclease-digested and concentrated virus was resuspended in the same Tris-EDTA buffer as above and treated with a mixture of dithiothreitol and EGTA at room temperature for 30 minutes to disrupt the chemical bonds holding the viral structural proteins together. After treatment with dithiothreitol and EGTA the sample was stored at minus 20° C for at least three hours. This treatment was repeated two more times. Following the three treatments, the virus preparation was centrifuged on a glycerol gradient as described for minichromosomes and the same fractions pooled as chromatin from disrupted virions.

### Chromatin Immunoprecipitation

We have performed our chromatin immunoprecipitation (ChIP) analyses according to the detailed description of the procedures which were published in 2017 with minor modifications as described below (Balakrishnan and Milavetz, 2017a, b). All of the antibodies used were ChIP validated by their respective vendors. Antibodies (and their vendors) included: hyperacetylated H3 (06-599, Millipore), hyperacetylated H4 (06-866, Millipore), H3K9me1 (ab9045, Abcam), H3K9me3 (ab8898, Abcam), and SP1 (39058, Active Motif). ChIPs were performed using Millipore kits according to the supplier’s protocol. In a standard ChIP, 10 µl of antibody (10 µg) was incubated with the chromatin being analyzed for 4 hours at 4^0^. A mixture of magnetic beads containing protein A and protein G was added (15 µl) magnetic beads and associated antibody bound chromatin was then purified according to the protocol present in the kit. In order to determine the percentage of the chromatin that was present bound to the chromatin, the bound chromatin was eluted using the buffer supplied in the kit and the DNA purified using Zymo Research ChIP DNA Clean and Concentrator columns according to the supplied protocol followed by elution in 26 μl H_2_O. The amount of DNA in the antibody bound fraction was then quantitated along with an aliquot of the input SV40 chromatin by real-time qPCR using primers that recognize the SV40 enhancer region.

ChIP-Seq was performed as above until the final elution step. Instead of eluting the intact SV40 chromatin bound to magnetic beads, the chromatin bound to beads was sonicated to fragment the chromatin and the fragments bound to the beads separated from the unbound fragments. The beads were washed twice with TE buffer from the kit followed by removal of the bound DNA using the elution buffer supplied in the kit. The DNA was again purified as above for preparation of sequencing libraries.

### Preparation of sequencing libraries from ChIP samples of SV40 chromatin

The DNA in the samples obtained from ChIP analyses was first purified using Zymo Research ChIP DNA Clean and Concentrator columns according to the supplied protocol and eluted in 26 μl H_2_O. In order to determine the percentage of SV40 chromatin immunoprecipitated, 1 µl was PCR amplified along with the starting chromatin used in the ChIP. The remaining DNA sample was dried prior to reconstitution with 25 µl of water during the preparation of the libraries. Libraries were prepared using the New England Biolabs Next UltraII kit according to the protocol supplied with the kit with one exception. All reagent volumes were reduced by half for the preparation of libraries with no apparent effect on library quality or quantity. Libraries were purified on Zymo Research ChIP DNA Clean and Concentrator columns according to the supplied protocol and eluted in 10 μl nuclease-free water. The libraries were then purified by submerged agarose gel electrophoresis using BioRad certified low-melting molecular biology grade agarose and a band corresponding in size to 200-300 base pairs of library DNA was cut out from the agarose gel. The agarose gel fragment containing the size-selected library DNA was purified on Zymo Research Gel DNA Recovery columns using the protocol and reagents in the kit and eluted from the columns in 21 μl nuclease-free water. The size-selected library fraction from each sample was amplified by PCR using indices from New England Biolabs and purified on AMPure as previously described (Balakrishnan and Milavetz, 2017a).

### Next Generation Sequencing (NGS)

Prior to sequencing, all of the libraries were analyzed for quality and quantity using an Agilent Bioanalyzer. Libraries which did not meet either the quality or quantity requirements were discarded. The libraries were sequenced on an Illumina MiSeq using protocols and reagents from Illumina in the epigenetics core laboratory at the University of North Dakota. For each combination of SV40 chromatin and antibody targeting a histone modification, a minimum of four libraries were prepared from ChIPs of distinct biological samples for sequencing. The actual number of libraries sequenced for each analysis is indicated in the figure legends. Preliminary quality control analysis of FastQ files was performed using FastQC v.0.11.2 (Andrews, 2010). The three prime adapters were trimmed from the reads using scythe v0.981 (Buffalo, 2011). Quality trimming was carried out using sickle v1.33 (Joshi and Fass, 2011) with a phred score of 33 as the quality threshold; reads with a length less than 45 bp were discarded. Reads were aligned to the SV40 genome (RefSeq Acc: NC_001669.1), cut at nucleotide (nt) 2666, using Bowtie-1.1.1 (Langmead et al., 2009) for wild-type SV40. Bedraphs were generated from aligned reads using Bedtools v2.23.0 (Quinlan and Hall, 2010). Before identifying peaks, fragments with a length greater than 200 bp were removed. Peak-calling was performed using the R/bioconductor package nucleR v2.20 (Flores and Orozco, 2011). Heatmaps were generated by merging the data from individual bedgraphs following normalization of the bedgraph.

### HIS-tagged SP1 binding to SV40 chromatin

HIS-tagged SP1 binding analyses were performed using reagents present in the Millipore ChIP kit. SV40 chromatin (25 μl) was incubated with an equal volume of chromatin dilution buffer from the kit, 2 μl of a 10 mg/ml solution of bovine serum albumin (Sigma), and 0.25 μg of HIS-tagged SP1 (Creative BioMart) for 30 minutes at 4^0^ with constant rotation. Ni-magnetic beads (15 μl) were added to bind the HIS-tagged SP1 and the mixture was incubated for an additional one hour with rotation. Following this incubation, the beads were washed according to the protocol in the ChIP kit as if the beads were derived from a ChIP analysis. The DNA present in the chromatin bound to the magnetic beads was eluted using the Kit buffer and purified using Zymo Research ChIP DNA Clean and Concentrator kits. The purified DNA was amplified by PCR as previously described and compared to a similar aliquot of the input chromatin to determine the percentage of the input chromatin bound by the HIS-tagged SP1.

HIS-tagged SP1-ChIP analyses were performed using a strategy similar to that described above for ChIP-ChIPs. SV40 chromatin was first bound by HIS-tagged SP1 as above and the bound chromatin purified. The bound chromatin was re-suspended in Tris-EDTA buffer (100 μl) and sonicated to fragment the chromatin. The bound fraction was separated from the unbound chromatin which was then added to a solution containing the antibody of interest (10 μl) and ChIP dilution buffer (400 μl). The fragmented chromatin was incubated with the antibody for 4 hours at which time protein A and G magnetic beads were added, followed by a further incubation for 4 hours. The chromatin fragments bound to magnetic beads were then purified using the Millipore ChIP kit protocol. The DNA associated with the beads was purified and amplified by real-time qPCR using primers that recognize the SV40 enhancer region in comparison to an aliquot of the chromatin fragment input in order to determine the percentage of the input chromatin which was bound by the antibody.

## ACKNOWLEDGEMENTS

The authors would like to thank Hannah Ness and the Epigenetic Core Laboratory at the University of North Dakota for sequencing the samples and the bioinformatics analyses. This work was funded by grants from the National Institutes of Health, AI094441 (to B.M.), AI142011 (to BM), GM104360 (to UND Epigenetics Core), and GM128729 (to Dakota Cancer Collaborative on Translational Activity).

